# Allopolyploidization from two dioecious ancestors leads to recurrent evolution of sex chromosomes and reversion to autosomes

**DOI:** 10.1101/2023.10.18.562857

**Authors:** Li He, Yuàn Wang, Yi Wang, Ren-Gang Zhang, Yuán Wang, Elvira Hörandl, Judith E. Mank, Ray Ming

## Abstract

Polyploidization presents an unusual challenge for species with sex chromosomes, as it can lead to complex combinations of sex chromosomes that disrupt reproductive development. This is particularly true for allopolyploidization between species with different sex chromosome systems. Here we assemble haplotype-resolved chromosome-level genomes of a female allotetraploid weeping willow (*Salix babylonica*) and a male diploid *Salix dunnii* using Hi-C and PacBio HiFi reads. We use phylogenomics of nuclear and plastid genomes to show that weeping willow arose from crosses between female ancestor from the *Salix* clade, having XY sex chromosomes on chromosome 7, with a male ancestor from the *Vetrix* clade, having ancestral XY sex chromosomes on chromosome 15. Our analysis reveals that weeping willow has one pair sex chromosomes, ZW on chromosome 15, that derive from the ancestral XY sex chromosomes in the *Vetrix-*clade male ancestor, and the X chromosomes on chromosome 7 from the *Salix-*clade female ancestor has reverted to an autosome. Taken together, our results point to rapid evolution and reversion of sex chromosomes following allopolyploidization in weeping willow.

**Significance Statement:** We assembled haplotype-resolved genomes and obtained gap-free sex chromosomes of a female allotetraploid weeping willow (*Salix babylonica*) and a male diploid *Salix dunnii*. The weeping willow arose from two dioecious ancestors, that have XY sex chromosomes on chromosome 7 and 15, respectively. The one pair sex chromosomes 15W and 15Z in weeping willow derived from ancestral 15X and 15Y, respectively. Inversions contributed to the evolutions of sex-linked regions (SLRs) of diploid and polyploid willows.

## Introduction

Polyploidy is an important feature of genome evolution (1, 2). Although highly degenerate sex chromosomes might act as a barrier to polyploidy (3), polyploidy in lineages with less differentiated and minimally degenerate sex chromosomes have been observed (e.g., dioecious plants, amphibians, and fishes, etc.) (1, 4, 5), and in some forms, may result in the rapid evolution of sex determining systems. An autopolyploid lineage, where both genome copies arise from the same species (6), could inherit the sex chromosomes of the diploid ancestor (7). However, allopolyploidy, where hybridization between species results in copies of each ancestor genome, could lead to complex sex chromosome combinations when the diploid ancestor species carry different sex chromosome types (XX/XY, male heterogametic and ZW/ZZ, female heterogametic), or even the same sex chromosome type but on different chromosomes (8, 9). This could lead to complex combinations of sex chromosomes that could disrupt sex determination process, leading to transitions between sex-determining systems, or even the loss of dioecy (10–12).

Most cases of polyploidization are followed by rediploidization, sometimes instantaneously from the point of polyploidization, but more often rediploidization is a gradual process (13). Rediploidization presents peculiar issues for duplicated sex chromosomes, as only one pair is retained as the sex determining chromosomes (4, 14), as has been observed in *Rumex acetosella* (15). Presumably the superfluous pair of sex chromosomes transition into autosomes, however the XY or ZW chromosomes of a pair may carry different gene content, and some genes might be lost in the rediploidization process (4). The retention and loss of sex chromosomes in allopolyploidization, and how that affects gene content, remains largely unexplored.

The role of polyploidization in other aspects of sex chromosome evolution is complex and not fully explored. For example, inversions on either the X or Y, or Z or W, can expand the non-recombining sex-linked region (16–20). It remains unclear how sex-linked inversions in polyploids differ, if at all, from diploids. Additionally, the sex-linked Y and W chromosomes often undergo genetic degeneration following recombination suppression (reviewed by (21–23)), resulting in gene dose differences between males and females. Dosage compensation mechanisms have evolved in some species that increase expression levels in the heterogametic sex, potentially restoring expression to levels in the ancestral chromosome pair, and making them equal in both sexes (24–27). However, polyploidization complicates the dose differences from Y and W chromosome degeneration, and the role of polyploidization in dosage compensation is largely unknown.

The weeping willow (*Salix babylonica*) is one of the best known trees in the world, and is an allotetraploid that possibly arose from offspring of crosses between species from clades *Salix* and *Vetrix* of genus *Salix* (9, 28, 29). Interestingly, the diploid species within the *Salix* clade have XY sex chromosomes on chromosome 7, whereas species in the *Vetrix* clade have XY or ZW sex chromosomes on chromosome 15 (30). The weeping willow is dioecious and produces viable seeds (28, 31), suggesting that it also has sex chromosomes and has overcome the allele dosage effect of the two different sex chromosome systems of its ancestors. This system provides an exciting opportunity to understand how allopolyploidization resulting from hybridization of two species with different sex chromosome systems resolves into stable dioecy.

The genetic control of sexes is in Salicaceae well known from poplars, the sister genus of *Salix*. In the male-specific region of the Y chromosome, duplicates of the partial ARABIDOPSIS RESPONSE REGULATOR 17 (*ARR17*) orthologue produce small RNAs and silence the intact *ARR17*-like gene of poplar sex chromosome 19, thereby suppressing female development (32). Partial *ARR17*-like duplicates were also found in Y-linked regions (7Y and 15Y) of willows, which can produce small RNA and suppress the expression of intact *ARR17*-like genes on chromosome 19 (33). The species of *Vetrix* clade inherited ancient sex-linked genes from their 15XY ancestor, ancient Y-linked genes is close to partial *ARR17*-like duplicates (30, 33, 34). There were two X-Y homologous gene pairs diverged in ancestor of *Vetrix* clade (34). These ancient sex-linked genes of *Vetrix* clade should be kept in weeping willow. We assembled a female haplotype-resolved chromosome-level genome of the allotetraploid weeping willow and a male haplotype-resolved genome of diploid *Salix dunnii* of the *Salix* clade with an XY system on chromosome 7, which has not previously been sequenced. We used our assemblies and other available genomes of willows to investigate the ancestors of the allotetraploid weeping willow. Our results suggest that the weeping willow inherited two alternative sex chromosomes, one on chromosome 7 from the female ancestor, and one on chromosome 15 from the male ancestor. Following polyploidization, the chromosome 7 sex chromosomes reverted to autosomal inheritance, while the chromosome 15 system transitioned from male to female heterogamety to form the system of stable dioecy observed in the weeping willow today.

## Results

### Genome assembly and annotation

From a single male *S. dunnii* individual (with XY sex chromosomes on chromosome 7 (35)), we generated 39 Gb (112X) of HiFi reads, 40 Gb (108X) of Illumina reads, and 37 Gb (107X) of Hi-C reads. From a single female *S. babylonica*, we generated 46 Gb (64X) of HiFi reads, 79 Gb (94X) of Illumina reads and 164 Gb (254X) of Hi-C reads (Table S1). After assembling each species separately with the PacBio HiFi and Hi-C reads, we used Illumina short reads to error-correct each genome. We obtained an *S. dunnii* assembly of 696 Mb, comprised of 40 contigs (contig N50 = 19 Mb) and a final gap-free chromosome-scale *S. dunnii* genome with 38 pseudochromosomes (Table1, Fig. S1). *S. babylonica* assembly was 1,286 Mb, comprised of 102 contigs (contig N50 = 16 Mb) (Table 1), with a final chromosome-scale assembly of 76 pseudochromosomes and gap-free sex chromosomes (Table S2, Fig. S2).

**Table 1.** Statistics of the *S. dunnii* and *S. babylonica* genome assemblies.

|  | *S. dunnii* | *S. babylonica* |
| --- | --- | --- |
| Total assembly size (Mb) | 696 | 1286 |
| Total number of contigs | 40 | 102 |
| Maximum contig length (Mb) | 37 | 32 |
| Minimum contig length (Kb) | 156 | 64 |
| Contig N50 length (Mb) | 19 | 16 |
| Contig N50 count | 16 | 33 |
| Contig N90 length (Mb) | 13 | 10 |
| Contig N90 count | 33 | 70 |
| Gap number | 0 | 24 |
| GC content (%) | 33 | 34 |
| Gene number | 65,485 | 123,231 |
| Repeat content (%) | 48 | 41 |

For the *S. dunnii* assembly, about 99.95% of Illumina short reads and 99.86% of HiFi reads were mapped back to the genome assembly, and around 99.18% and 99.86% of the assembly was covered by at least 20× reads, respectively. Similarly, about 98.91% of Illumina short reads and 99.43% HiFi reads could be aligned back to the *S. babylonica* genome assembly, and 99.41% and 95.79% of the assembly was covered by at least 20× reads, respectively. BUSCO analysis suggested that 1416 (98.4%) highly conserved core proteins in the *S. dunnii* genome, and 1422 (98.8%) highly conserved core proteins were identified in the *S. babylonica* genome (Table S3).

We recovered 333.7 Mb (47.92%) and 525.5 Mb (40.86%) of repetitive sequences in the *S. dunnii* and *S. babylonica* genomes (Table S4). Among them, the percentage of LTR was the highest, accounting for 24.78% in *S. dunnii* and 19.37% of *S. babylonica* (Table S4). A total of 65,485 genes, (61,872 mRNAs excluding alternative splicing, 1,139 transfer RNAs (tRNAs), 1,102 ribosomal RNAs (rRNAs) and 1,372 unclassifiable noncoding RNAs (ncRNAs)) were identified in *S. dunnii* (Table S5). Similarly, we obtained 123,231 genes (116,745 mRNAs, 2,428 tRNAs, 1,528 rRNAs and 2,530 ncRNAs) in *S. babylonica* (Table S6). The average *S. dunnii* gene is 3683.4 bp long including 6.4 exons, and the average *S. babylonica* gene is 3421.6 bp long including 5.9 exons (Table S7). By combining several strategies, we were able to match the vast majority of our predicted genes to predict protein in public databases such as GO, KEGG, and Swiss_Prot (Table S8), and only 2.24% *S. dunnii* protein-coding genes and 1.77% of *S. babylonica* protein-coding genes were not able to be annotated.

### Phylogenetic analysis and divergence time estimation

We found that *S. babylonica* is tetraploid based on flow cytometry and allotetraploid according to *k*-mer results (Fig. S3 & Fig. S4). To determine the position and origin of the two subgenomes within the genus *Salix*, we obtained 5,146 single-copy orthologs from our *S. babylonica* (Aa and Ba haplotypes) and *S. dunnii* (a haplotype) genome assemblies, as well as six other willows with available genomes, and the outgroup *Populus trichocarpa* (Table S9). The resulting species tree is broadly consistent with previous studies and separated the *Salix* and *Vetrix* clades (33–36). Importantly, our results show that the *S. babylonica* B subgenome originated from an ancestor in the *Vetrix* clade and diverged from the most recent common ancestor about 24 Mya (million years ago), while the A subgenome originated from an ancestor in the *Salix* clade, diverging from the most recent common ancestor about 20.95 Mya (Fig. 1(a)). We also built a chloroplast genome phylogeny, which suggests that the maternal ancestor of *S. babylonica* originated from the *Salix* clade (Fig. 1(b), S5).

**Figure 1.**
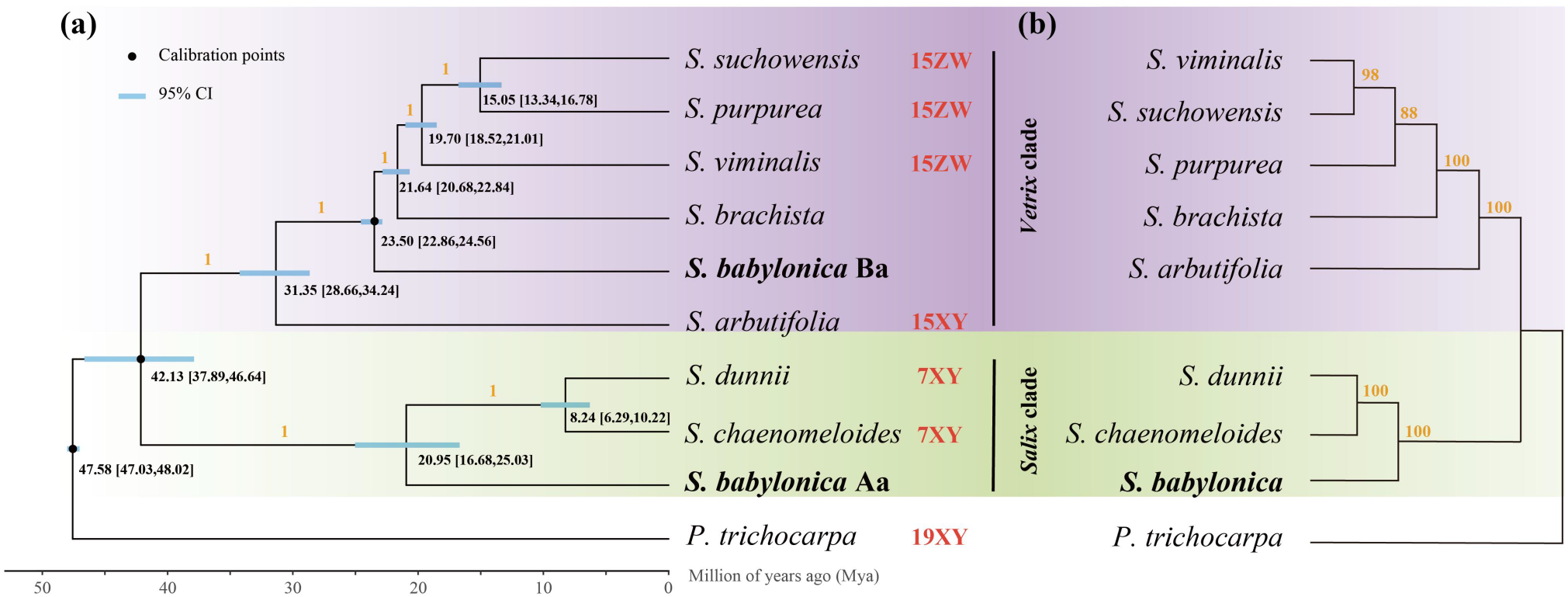
Origin and divergence time estimation of *S. babylonica* subgenomes. (a) Inferred phylogenetic tree and divergence times based on genome sequences of eight willows and the outgroup *P. trichocarpa*. Blue node bars are 95% confidence intervals, and the black nodes are three fossil calibration points (36). Black numbers marked around nodes represent divergence time. Orange numbers marked above branches represent support values. Red characters on the right represent different sex determination systems. (b) Inferred phylogenetic tree based on chloroplast genome sequences of the nine species. Numbers marked on the tree represent bootstrap values (Fig. S5).

According to the inferred time of transposable elements divergence (see methods in detail) between subgenomes of *S. babylonica*, the estimated allotetraploidization time is around 5.85–2.75 Mya (Fig. S6). Therefore, the *S. babylonica* may have emerged during 5.85–2.75 Mya, suggesting allotetraploidization in this species without human influence.

### Identification of the sex determination system

We used 4,417.1 million clean Illumina reads from 20 females and 20 males of weeping willow (97 - 135.8 million reads per individual, mean 110.4, Table S10), to identify sex-specific-*k-*mers, and found a significant enrichment of female specific *k*-mers on chromosome 15Ba, between 5.32–9.71 Mb (Fig. 2(a), Fig. S7), consistent with a W chromosome. Our results from the chromosome quotient (CQ) method (37) are concordant with the *k*-mer analysis, and we detected the sex-linked region between 5.28–9.69 Mb on chromosome 15Ba, with mean M:F (male : female) CQ 0.27, consistent with a W chromosome. The M:F CQ of 2.83–7.88 Mb on chromosome 15Bb had a mean of 2.1, consistent with a Z chromosome. These results suggest that weeping willow has a female heterogametic system on chromosome 15, 15Ba and 15 Bb are W and Z chromosomes, respectively. Our synteny analysis revealed two inversion events between the W and Z chromosomes of *S. bablyonica*, and further confirmed a 6.3 Mb W-SLR region between 3.41–9.71 Mb on 15Ba and 5.26 Mb Z-SLR counterpart between 2.66–7.92 Mb on 15Bb (Fig. 2(a), Fig. S8). The other regions of the two chromosomes form pseudoautosomal regions (PARs) with 8.74 Mb in 15Ba and 8.12Mb in Bb, respectively.

**Figure 2.**
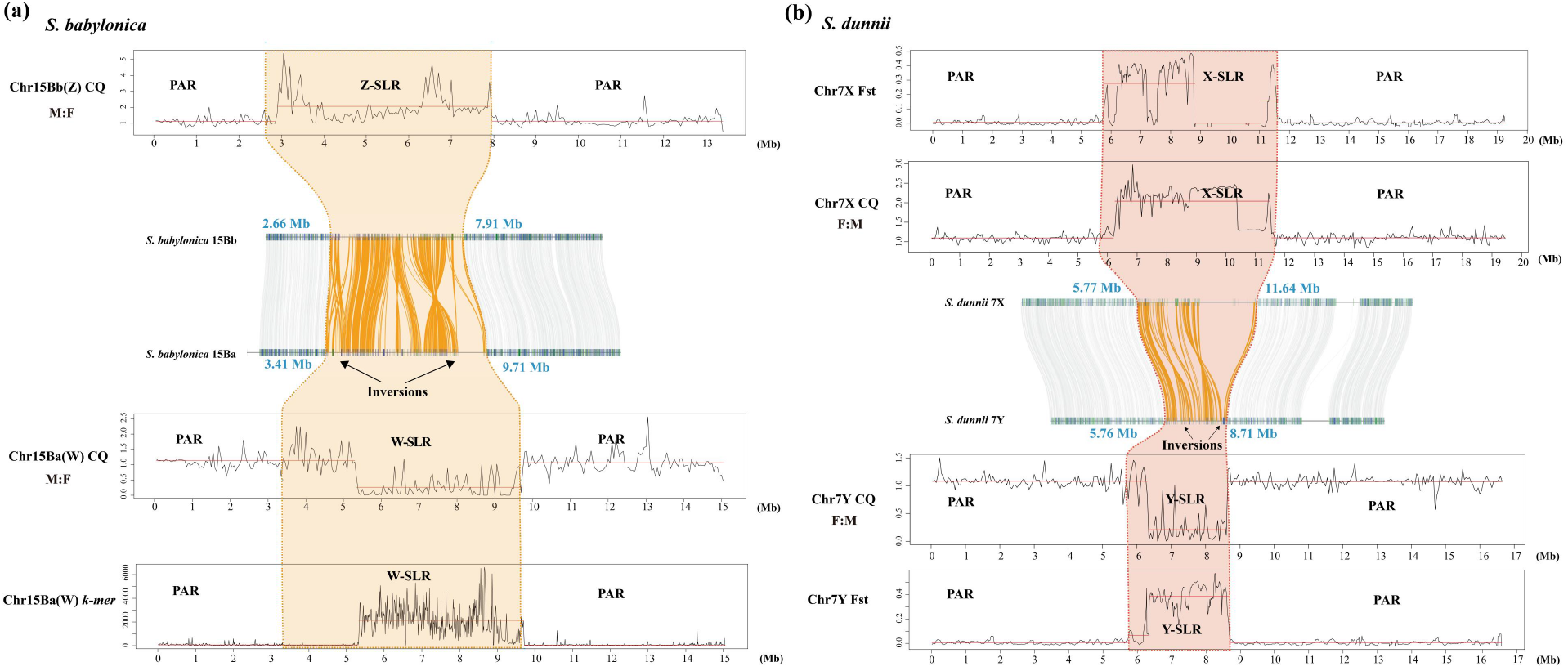
Sex-linked regions in *S. babylonica* and *S. dunnii*. (a) The CQ results (male vs. female alignments) on sex chromosomes 15Bb(Z) and 15Ba(W) of *S. babylonica*, and the female specific *k-mer* signals on chromosome 15Ba. Fig. S7 presented the the *k-mer* results of whole genome. The collinearity results between two sex chromosomes show two inversions. (b) The *F*st results between the sex groups on sex chromosomes 7X and 7Y of *S. dunnii*, and the CQ results (female vs. male) on sex chromosomes 7X and 7Y. The collinearity results between two sex chromosomes show three inversions.

The *S. dunnii* has a male heterogametic system on chromosome 7 (35). The F:M (female : male) CQ analysis revealed a 5.35 Mb region, 6.15–11.5 Mb, on 7X with mean CQ 2.03, and a 2.35 Mb region, 6.25–8.6 Mb, on 7Y with mean CQ 0.22 (Fig.2(b)). We obtained 6,687,746 and 6,705,953 SNPs using a (including 7X) and b (including 7Y) haplotype genomes of *S. dunnii* as references, and calculated the *F*_ST_ values between the two sex groups based on the SNPs. Changepoint analyses detected significantly higher *F*_ST_ values between 5.77–11.64 Mb on chromosome 7X and 5.76–8.71 Mb on chromosome 7Y than in other regions (PARs), which covered the regions identified by CQ. We identified three inversion events between the X and Y in *S. dunnii* using synteny analysis (Fig. 2(b), Fig. S9), which are all within the region identified as sex-linked by *F*_ST_ analysis.

### Evolutionary origin of *Salix babylonica*

The nuclear and chloroplast trees imply that the female ancestor of *S. babylonica* most likely arose from the *Salix* clade, while the male ancestor most likely came from the *Vetrix* clade. The *Vetrix* clade contains both 15XY and 15ZW species, however phylogenetic analysis of ancient sex-linked genes within the *Vetrix* clade, the X and -Y linked genes diverged in ancestor of the *Vetrix* clade (34), indicating that the 15XY is the most likely sex chromosome system for the *S. bablyonica* male *Vetrix* ancestor (Figs. 1, 3, S10). We assume here a scenario of two diploid ancestors and both forming unreduced gametes, having chromosome doubling after fertilization, forming allopolyploids via a triploid bridge, or crosses between two tetraploid ancestors would also be possible (6, 38). This suggests that the unreduced gametes from the male *Vetrix* ancestor gametes were likely 15X15Y (skipping meiosis I), or 15X15X and 15Y15Y (skipping meiosis II), and from the female *Salix* ancestor gametes were likely 7X7X. These female and male gametes can produce 7X7X15X15Y (now 7Aa7Ab15W15Z) female and 7X7X15Y15Y (now 7Aa7Ab15Z15Z) male.

**Figure 3.**
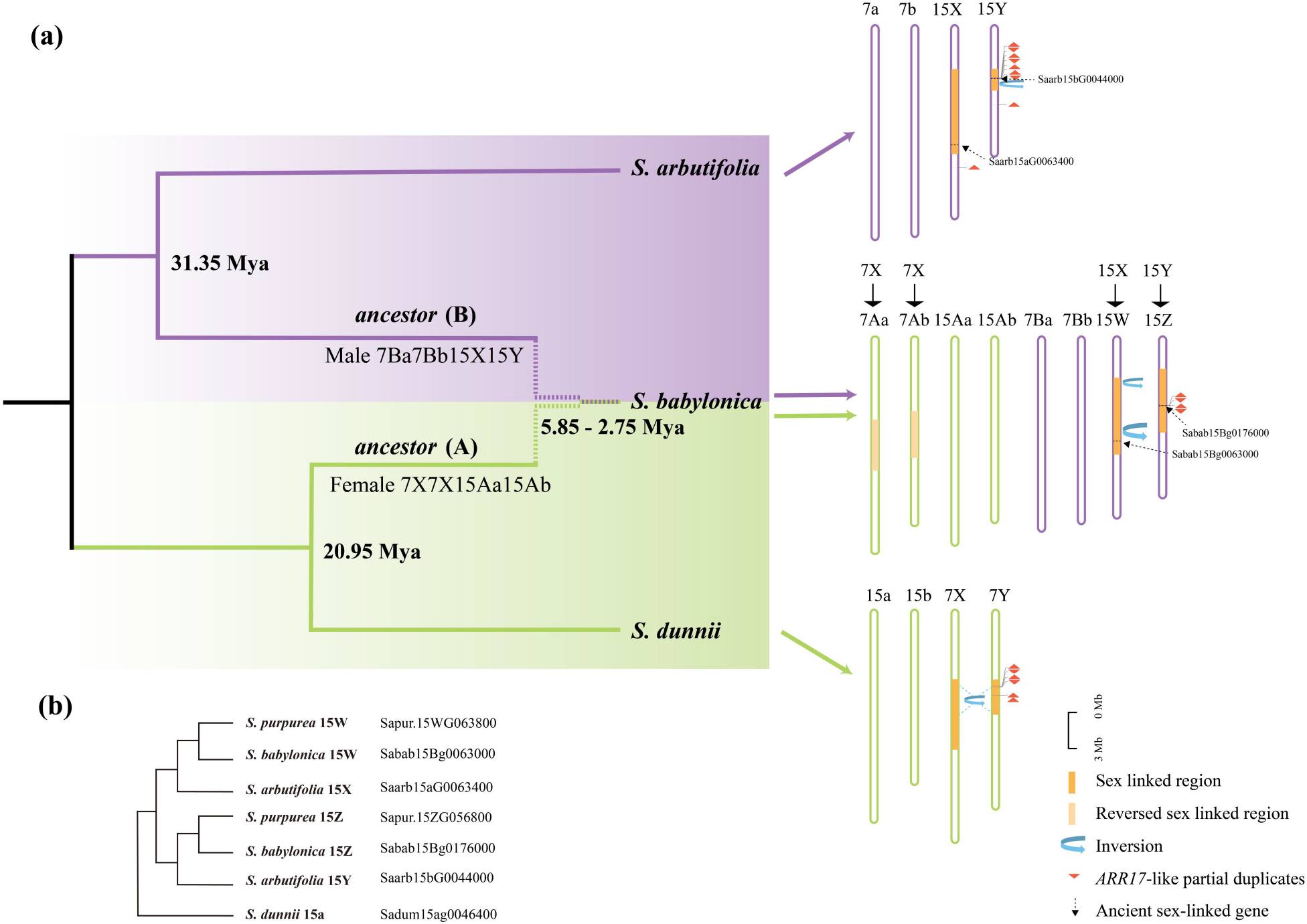
Hypothetical origination of *S. babylonica* and evolution of sex-linked regions in relevant diploid willows. (a) The purple background represents genomic contents of *S. babylonica* from ancestor of the *Vetrix* clade, that likely has 15XY sex determination system (SDS), while the green background shows the genomic components from ancestor of the *Salix* clade, that has 7XY SDS. The chromosomes on the right use the scaled real physical length, and dark yellow or light yellow on them marked the sex-linked or reversed sex-linked regions (Fig. S12). The red triangles within sex-linked regions mean *ARR17*-like partial duplicates (pointing down represents forward), and blue arrows shows the inversions emerging on SLRs. The genes marked by the black arrows are ancient sex-linked genes that used for (b). (b) The phylogenetic tree is reconstructed from ancient sex-linked genes that proposed by (34) (Fig. S10).

To better understand the transition from 15XY in the *S. babylonica* male *Vetrix* ancestor to the current *S. babylonica* 15ZW sex chromosome system, we examined the distribution of *ARR17*, which is involved in sex determination in *Salix* and *Populus* (32, 33), in *S. arbutifolia* (15XY) and *S. babylonica*. We found *ARR17-*like partial duplicates on 15Y of *S. arbutifolia* and 15Z of *S. babylonica* (Fig. S11), while they are absent in 15X and 15W of these two species. This further suggests that the ancestral *Vetrix* 15Y transitioned to 15Z in *S. bablyonica,* and 15X transitioned to 15W.

We also assessed collinearity between sex chromosome (7X and 7Y) of *S. dunnii* and homologous autosome (7Aa) of *S. babylonica*. Our results suggest that 7X of female *Salix* ancestor has reverted to autosomal inheritance in *S. bablyonica* (Fig. S12). There are inversions between sex chromosomes of *S. dunnii*, but the gene order in autosome 7Aa of *S. babylonica* is most similar to 7X of *S. dunnii*, again consistent with a female *Salix* ancestor (Fig. 3(a) and Fig. S13).

### Evolution of the sex-linked region

Our analysis reveals *ARR17*-like partial duplicates located in the inversions between 7X and 7Y of *S. dunnii*, and close the *S. babylonica* inversion between 15Z and 15W and the *S. arbutifolia* inversion between 15X and 15Y (Fig. 3(a), Table S11). It is difficult to determine whether the detected inversions are a catalyst or consequence of recombination suppression (39), however these results suggest that *ARR17*-like partial duplicates are associated with sex-linked inversions. This is either because recombination has been selectively suppressed in these regions to maintain sex-specific segregation patterns of the *ARR17*-like partial duplicates, or alternatively the inversions may have resulted from the fact that these loci are in areas of pericentromeric region with low recombination (Fig. S14, Fig. S15, (34)). We further compared the expansion of the *S. babyloncia* and *S. arbutifolia* sex-specific regions (Fig. S16), and our results indicated that these regions evolved independently in different sex chromosomes despite having the same origin.

In *S. dunnii* sex-linked region, including 5.87 Mb X-SLR and 2.95 Mb Y-SLR, this is consistent with *S. arbutifolia* (7.06 Mb X-SLR and 1.81 Mb Y-SLR) (34), but it is opposed to *Silene latifolia*, which has longer Y-SLR than X-SLR(19, 40). There are 128 and 96 protein coding genes within X-SLR and Y-SLR in *S. dunnii* (Table S5), and the X-SLR has 2.72 Mb longer total repeated sequences than the Y-SLR (Table S12). We also identified 31 (13 of them are tandem duplications) and 16 (3 of them are tandem duplications) specific genes in X-SLR and Y-SLR, respectively (Table S13). Therefore, the accumulation of repeated sequences and specific tandem duplications may contribute to the longer X-SLR than Y-SLR in *S. dunnii*. This result suggests that *S. dunnii* with 7XY have specific sex chromosome evolution pattern compared the other plants with XY SDSs (41, 42).

### Dosage compensation in different sex chromosomes

We found degeneration signals in the sex-linked regions of the *S. babylonica* 15Z and 15W, *S. arbutifolia* 15X and 15Y, and *S. dunnii* 7X and 7Y. Gene density was lower in all these sex-linked regions compared to the corresponding PARs, while repetitive elements density was higher (Fig. S17; Table S12). The W-SLR and Y-SLRs had lost genes compared with their orthologous autosomes, i.e., *S. babylonica* 15W-SLR lost 2.84% genes, *S. arbutifolia* 15Y-SLR lost 10.59% genes, and *S. dunnii* 7Y-SLR lost 1.22% genes in (Table S14). We also found Z-SLR and X-SLRs had lost genes, i.e., *S. babylonica* 15Z-SLR lost 3.59% genes in), *S. arbutifolia* 15X-SLR lost 8.24% genes, and *S. dunnii* 7X-SLR lost 1.22% genes, which is consistent with (43). Z-SLR lost more genes than W-SLR in *S. babylonica*, which may be due to the turnover (Y→Z) mentioned above.

Overall, the expression values between sexes are similar but significantly lower than in autosomes among 15ZW (male 15Z-SLR15Z-SLR vs female 15Z-SLR), 15XY (female 15X-SLR15X-SLR vs male 15X-SLR) and 7XY (female 7X-SLR7X-SLR vs male 7X-SLR) (Fig. 4). Based on these results, we revealed similar incomplete dosage compensation pattern in *S. babylonica*, *S. arbutifolia* and *S. dunnii*.

**Figure 4.**
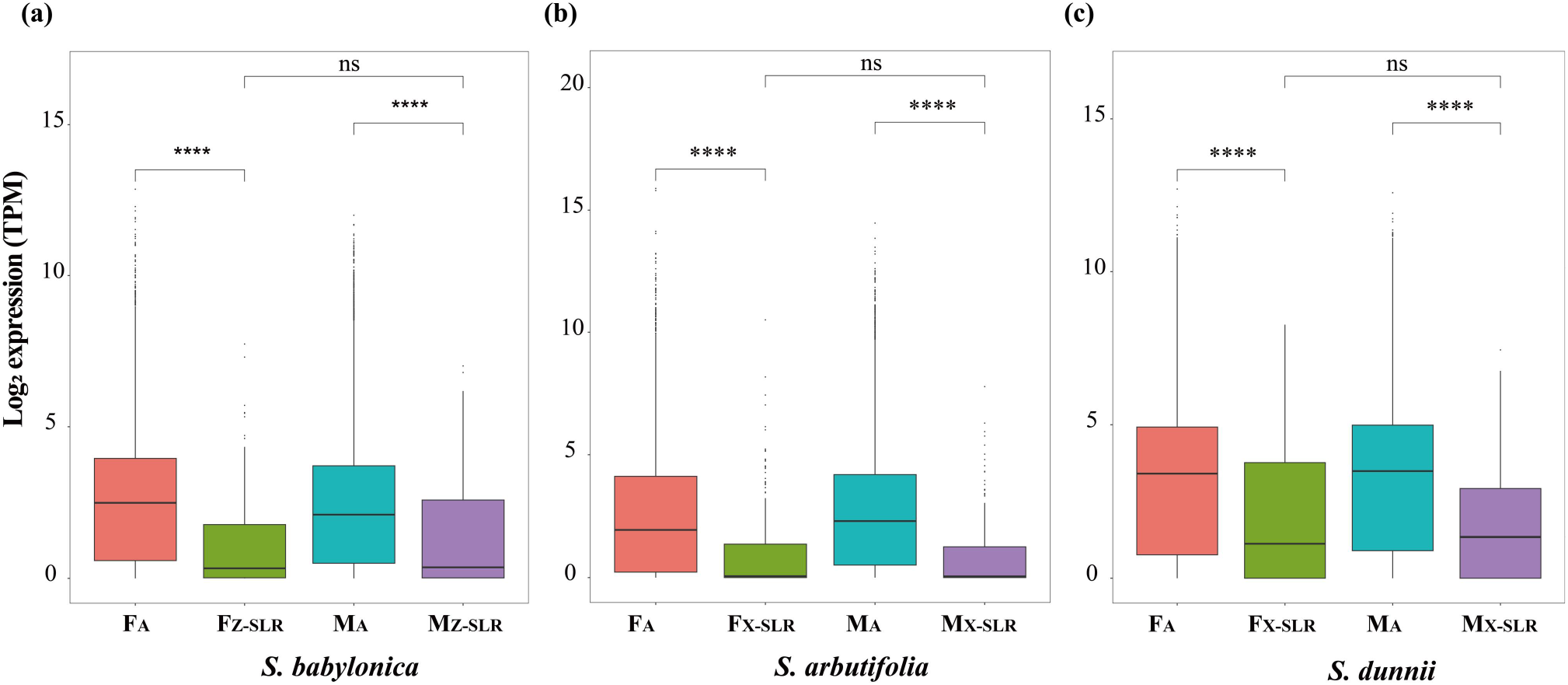
Dosage compensation pattern in different sex-linked regions of willows. The expression of genes in two sexes of willows and their autosomes (Wilcox test \**p* < .05) in *S. babylonica* (a), *S. arbutifolia* (b), and *S. dunnii* (c). F and M represent female and male. A and SLR represent autosome and sex-linked region.

## Discussion

Although sex chromosomes were once thought to be an obstacle to polyploidization (3), it is increasingly clear that polyploidy can arise from diploid ancestors with sex chromosomes in both plants and animals (4, 8). However, how these polyploid lineages overcome the dosage effect of polyploidization to establish stable dioecy is not yet known. Previous studies have not provided direct evidence that dioecious plants can break through the polyploidy limit. *Diospyros kaki* and *Mercurialis annua*, both male heterogametic, reverted to non-dioecious sex determination systems after genome doubling (44, 45), while the ancestors of the octoploid dioecious *Fragaria chiloensis* were hermaphroditic (46). For other polyploids with sex chromosomes, such as *Rumex acetosella*, a male heterogamety tetraploid, genomic resources were not available to investigate sex chromosome evolution and its ancestral species (15).

We assembled haplotype-resolved genomes of female allotetraploid *S. babylonica* and male diploid *S. dunnii*. Our nuclear and chloroplast genome sequences trees indicate that *S. babylonica* has a female ancestor from the *Salix* clade and a male ancestor from the *Vetrix* clade within the *Salix* genus and that allopolyploidization occurred between 5.85–2.75 Mya, long before plant domestication (47). Similar to its male ancestor, the *S. babylonica* sex-linked regions are located on chromosome 15Ba(15W) and 15Bb(15Z) according to *k-*mer and CQ analysis (Fig. 2(a)), indicating only one pair sex chromosomes were retained, as hypothesized by (14). According to the ancient sex-linked gene tree, that X-Y homologous gene pair diverged in ancestor of the *Vetrix* clade (34), the 15W(15Ba) and 15Z(15Ba) derived from the ancestral 15X and 15Y chromosomes in the male ancestor, respectively (Fig. 3; Fig. S11).

Early generations of allotetraploid *S. babylonica*, comprised probably 7X7X15X15X and 7X7X15X15Y females (see below), 7X7X15Y15Y males. Initially, the two 15Ys in males can restore the recombination rate, but cannot be inherited unless their gametes cross with a female gamete carrying 15Y, which can only be produced by a female-fertile 7X7X15X15Y genotype (Figure S13). Furthermore, the 7X7X15X15X can only produce 7X15X and crosses with a male gamete carrying 7X15Y, and produced 7X7X15X15Y genotype. In this process, recombination reduced in 15X of female, and which then transitioned to W. Our evidence suggests that the 7X chromosomes from the female ancestor reversed to autosomes, and share inversions with the X chromosome of *S. dunnii* (Fig. S12).

*ARR17*-like gene (partial and intact) acts as sex determination factor in some members of Salicaceae (32, 33). The partial *ARR17*-like duplicates in SLRs of 15Y can produce small RNA and silence intact *ARR17*-like genes on two chromosomes 19s (32, 33). In 7XY and 15XY SDSs, previous studies show the correspondence between one Y chromosome with partial *ARR17*-like genes and four intact *ARR17*-like genes in two Chr19s (Table S15, Figure S18 (33)). Diploid *S. chaenomeloides* (7XY) and *S. arbutifolia* (15XY) perform this pattern (33, 34), suggesting suppression effects between one Y chromosome with partial *ARR17*-like genes and four intact *ARR17*-like genes, hence determining male. The fact that partial *ARR17*-like duplicates are within/close inversions in the *S. babylonica* and *S. dunnii* sex chromosomes suggests that these loci act as sex determining factors in these species as well. Although it is unclear whether inversions are the cause of recombination suppression between sex chromosomes or/and are a consequence of it (39), the association between *ARR17-*like partial duplicates and suggests selection to suppress recombination in these regions to maintain sex-specific segregation patterns of the *ARR17*-like partial duplicates. The Z chromosome from Y chromosome can also suppress the intact *ARR17*-like genes in Chr19s (33, 34). However, in *S. purpurea* (15ZW), the four redundant intact *ARR17*-like genes in chr15W can still determine female although the four intact *ARR17-*like genes on two Chr19s may be suppressed by partial *ARR17*-like genes on Chr15Z (33). But *S. babylonica* likely achieved female sex determination by increasing the number of intact *ARR17*-like genes on four Chr19s (eight intact *ARR17*-like genes) (Figure S18).

*S. babylonica* and *S. arbutifolia* shared common ancestral 15X and 15Y, and our phylogenetic analysis of *ARR17-*like partial duplicates (Fig. S11) and ancient sex-linked orthologous genes (Fig. 3(b), (34)) suggest that the ancestral *Vetrix* 15Y transitioned to 15Z in *S. bablyonica,* and 15X transitioned to 15W. However, we observe inversions on *S. babylonica*’s 15W, but the inversions on the *S. arbutifolia* 15Y. The sex-linked regions of *S. dunnii* have more complicated mutations, and we identified inversions in both 7X and 7Y. However, all these sex-linked regions likely overlap with the centromeric region, and it is therefore not possible to exclude the possibility that the low recombination of centromeric regions played a role in the formation of these inversions (34, 35).

Gene loss in sex-linked regions of sex chromosomes is common in plants (40). Although dioecy is ancient in willows, and originated at least 47 Mya, sex chromosomes in other species within the *Vetrix* clade, including *S. viminalis* (15ZW) (48) and *S. purpurea* (15ZW) (49) show low levels of divergence (35, 48). The results showed that 15Y-SLR of *S. arbutifolia* lost more genes compared with 15X-SLR (Table S14). This signature was also found in other plant sex chromosome systems, including papaya (50), *Silene latifolia* (40) and *Rumex hastatulus* (51) and is also a general feature of animal sex chromosomes (52). Our results suggested that the 15W in *S. babylonica* came from 15X, and 15Y came from 15Z. It is likely that *S. baylonica* inherited ancestor(B)’s characteristics including gene loss and expression, losing more genes in Z-SLR and performing similar incomplete dosage compensation pattern (Fig. 4). This pattern was also found in *S. dunnii* (7XY).

## Materials and Methods

### Plant material

We collected young leaves from a male *Salix dunnii* (FNU-M-1) and female weeping willow *S. babylonica* (saba01F) plant for genome sequencing. Young leaf, catkin, stem and root samples for transcriptome sequencing were collected from FNU-M-1 and saba01F, as well as catkins from two other female and three male plants of weeping willow, one other female plant of *S. dunnii*. The plant material was frozen in liquid nitrogen and stored at −80°C until total genomic DNA or RNA extraction. We sampled 40 individuals of *S. babylonica* and dried their leaves in silica-gel for resequencing. Voucher specimens collected for this study are deposited in the herbarium of Shanghai Chenshan Botanical Garden (CSH). Table S16 gives detailed information of all the samples. We also downloaded genome sequence data of 38 individuals of *S. dunnii* published by (35), the genome of *S. arbutifolia* ((34), with gap-free 15X and 15Y sex chromosomes), and other available willows genomes and *Populus trichocarpa* for relevant analysis (Table S9).

### Ploidy determination

The ploidy of saba01F was measured by flow cytometry, using diploid *Salix dunnii* (2x = 2n = 38, (35)) as external standard. The assay followed the protocol of (53). The leaf tissue was incubated for 80 min in 1 mL LB01 buffer and chopped with a razor blade. The homogenate was then filtered through a 38-μm nylon mesh and treated with 80 μg/mL propidium iodide (PI) and 80 μg/mL RNase followed by 30 min incubation on ice to stain the nuclei. DNA content measurements were done in MoFlo-XDP flow cytometer (Beckman Coulter Inc.) and evaluated using FloMax version 2.0 (Sysmex Partec GmbH). The ploidy level was calculated as sample ploidy = reference ploidy × mean position of the sample peak/mean position of reference peak.

### Determination of allo- or autotetraploid origin in weeping willow

We used jellyfish version 2.3.0 (54) to construct *k*-mer frequency distributions of saba01F based on PCR-free Illumina short reads, with *k*-mer length set to 21. Genomescope 2.0 (55) was run with 21-mer to distinguish between autotetraploid and allotetraploid origin based on the patterns of nucleotide heterozygosity. Allotetraploids are expected to have a higher proportion of aabb than aaab, while the autotetraploids have a higher proportion of aaab (7, 55).

### Genome sequencing

For Illumina PCR-free sequencing of FNU-M-1 and saba01F, and whole-genome sequencing (WGS) of all 40 weeping willow individuals, their total genomic DNAs were extracted using Qiagen DNeasy Plant Mini kit following the manufacturer’s instructions (Qiagen). PCR-free sequencing libraries were generated using the Illumina TruSeq DNA PCR-Free Library Preparation Kit (Illumina) following the manufacturer’s recommendations. Paired-end libraries were constructed for all 40 samples for WGS. These libraries were sequenced on an Illumina platform (NovaSeq 6000) by Beijing Novogene Bioinformatics Technology (hereafter Novogene).

The Hi-C library was prepared following standard procedures (56). The leaves from FNU-M-1 and saba01F were fixed with a 4% formaldehyde solution. Subsequently, cross-linked DNA was isolated from nuclei. The restriction enzyme MboI was then used to digest the DNA, and the digested fragments were labelled with biotin, purified and ligated before sequencing. Hi-C libraries were controlled for quality and sequenced on NovaSeq 6000 by Novogene.

For PacBio libraries and sequencing, total genomic DNA was extracted using the CTAB method. The genomic DNA was checked by agarose gel electrophoresis, showing a main band above 30 kb. PacBio large insert libraries were prepared with SMRTbell Express Template Prep Kit 2.0. These libraries were sequenced by Novogene using the PacBio Sequel II platform.

### RNA extraction and library preparation

Total RNA was extracted from young leaves, female and male catkins, stems and roots of *S. dunnii* and weeping willow using CTAB method. RNA integrity was assessed using the Fragment Analyzer 5400 (Agilent Technologies, CA, USA). Libraries were generated using NEBNext® UltraTM RNA Library Prep Kit for Illumina® (NEB, USA) following manufacturer’s recommendations, and sequencing was performed on an Illumina Novaseq 6000 by Novogene.

### Genome assembly

Hifiasm (57) was used to assemble initial genomes based on PacBio HiFi reads, and only the haplotype assemblies were used for subsequent analysis. For chromosome-level genome assembly, the Hi-C reads were first aligned to the haplotype contig genome using Juicer (58). Subsequently, a preliminary Hi-C-assisted chromosome assembly was carried out using 3d-dna (59). This was followed by manual inspection and adjustment using Juicebox (60), primarily focusing on refining chromosome boundaries, removing incorrect insertions, adjusting orientations and rectifying assembly errors. To optimize the assembly further, gap filling based on HiFi reads was performed using the LR_Gapcloser (61). Additionally, a two-round polishing approach using Nextpolish (62) was employed for genome base correction utilizing short-read data.

We obtained chromosome-scale, haplotype-resolved genome assemblies of *S. babylonica* and *S. dunnii*. Although the majority of chromosomes exhibited well-assembled telomeric ends containing the characteristic telomere sequence (TTTAGGG)n, there were a few cases where this sequence was either short or missing. Assuming incomplete assembly or insufficient extension, the HiFi reads were remapped to the genome. Reads aligning near the telomeres were selected, and contig assembly was performed using hifiasm (57). The resulting contigs were then aligned back to the chromosomes, allowing for extension of the chromosomes towards the outer ends to enhance the completeness of the telomere sequence assembly. For the *S. babylonica* genome assembly, the chromosomes were compiled as chromosome 01 to chromosome 19 (Aa, Ab, Ba, Bb) according to the homologous relationship with *S. dunnii* (A) and *S. brachista* (B). The chromosome number of our male *S. dunnii* is consistent with that of the female *S. dunnii* genome reported previously (35).

The chloroplast and mitochondrial genomes were assembled using GetOrganelle (63). Afterwards, fragmented contigs were mapped to the chromosome-level genome and organelle genome sequences by Redundans (64), enabling the identification of redundant segments. Furthermore, low-coverage fragments or haplotigs within the scattered sequences and rDNA fragments were discarded.

### Genome annotation

Homology-based prediction, transcript prediction, and *de novo* prediction approaches were used for genome annotation. For homology-based prediction, we used the publicly available Salicaceae protein sequences (Table S9) as homologous protein evidence for gene annotation. For transcript prediction, we used three strategies to assemble transcripts. In addition to using Trinity (65) to *de novo* assembly directly, the reads were also aligned to genomes with HISAT2 (66) before assembling with Trinity genome-guild model and StingTie (67). Then we combined all the transcript sequences and removed redundancy using CD-HIT (68) (identity > 95%, coverage > 95%). Based on transcript evidence, we used the PASA pipeline (69) to annotate gene structure, and aligned to the reference protein (Table S9) to identify full-length genes. These genes were used for AUGUSTUS (70) training, and we performed five rounds of optimization.

MAKER2 (71) was used for genome annotation based on *de novo* prediction, transcript and homolog protein evidence. After masking repeat sequences with RepeatMasker, AUGUSTUS (72) was used for *ab initio* predict coding genes. We then aligned transcript and protein sequences to the genomes using BLASTN and TBLASTX. After optimizing the comparison results using Exonerate (73), we integrated and predicted gene models with AUGUSTUS (72), and expression sequence tag (EST). EVidenceModeler (EVM) gene structure annotation tool (74) was further used to integrate MAKER and PASA annotation results. In addition, we performed TEsorter (75) and EVM to identity and mask protein domains. Then, PASA (69) was used to upgrade the EVM annotation, adding untranslated region (UTR) and alternative splicing. Finally, the abnormal coding frame (internal stop codon or ambiguous base, no start codon or stop codon) and too short (<50 amino acids) gene annotations were removed.

For non-coding RNAs (ncRNAs) annotation, tRNAs were annotated using tRNAScan-SE (76), and RfamScan was used to align and annotate various ncRNAs. We also performed Barrnap (https://github.com/tseemann/barrnap) to remove partial results. Finally, we removed the redundancies and integrated all the annotation results.

The functions of protein-coding genes were annotated based on three strategies. (1) Gene functions were identified using eggNOG-mapper (77), aligning with homologous gene databases; (2) We used DIAMOND to perform sequence similarity searches, and compared protein sequences with protein databases including Swiss_Prot, TrEMBL and NR (identity > 30%, E-value < 1e-5); (3) InterProScan (78) was used to search domain similarity. We compared sequences with the PRINTS, Pfam, SMART, PANTHER and CDD databases and obtained amino acid conserved sequences, motifs and domains.

To identify repetitive sequences in genomes, we used EDTA (79) to identify transposable elements (--sensive 1, --anno 1), producing a TE library. Then, RepeatMasker (http://www.repeatmasker.org/Robtained/) was used to determine repetitive regions within our genome assemblies.

### Phylogenetic analysis and divergence time estimation

We performed a phylogenetic analysis of the eight willow genomes (*Salix suchowensis*, *S. purpurea*, *S. viminalis*, *S. brachista*, *S. babylonica* (haplotypes Aa and Ba), *S. arbutifolia* (haplotype a), *S. dunnii* (haplotype a), *S. chaenomeloides*), and used *Populus trichocarpa* as outgroup (Table S9). After obtaining longest mRNAs among them, OrthoFinder (80) was used to identify single-copy orthologous genes. To avoid the effect of sex chromosomes, we removed genes from chromosomes 7, 15 and 19. Then, these protein sequences were aligned using MAFFT (81). We constructed gene trees with IQ-TREE (-m MFP -bb 1000 -bnni -redo) (82). Then, ASTRAL was used to infer a species tree based on gene trees results (83). Finally, MCMCTREE in PAML(84) was used to estimate the divergent time based on two fossils and one inferred time at three nodes: (1) the root node of Salicaceae (48 Mya), (2) the divergence time between *Salix* and *Vetrix* (37.15 Mya to 48.42 Mya), and (3) the ingroup of *Vetrix* (*Chamaetia*-*Vetrix*) clade (23 Mya) (36). The chloroplast genomes of the nine species were obtained from available assemblies or assembled using GetOrganelle (63) based on whole genome sequencing reads (Table S9). These chloroplast genome sequences were aligned by HomBlocks.pl with default parameter (85), and then IQ-TREE was used to construct chloroplast phylogenetic tree.

In order to determine the allotetraploidization time of *S. babylonica*, we calculated TEs (transposable elements) divergence values from the two subgenomes. This estimation method was described by (86), because TEs evolutionary rates of two subgenomes of allotetraploids different before and after polyploid formation, and therefore TEs substitutions of them also changed significantly during the polyplodization. Among the four haplotypes of *S. babylonica*, we used Aa vs. Ba and Aa vs. Bb to perform this analysis, removing chromosomes 7 and 15. First, we used Nucmer (87) to identify the matching region between two subgenomes of *S. babylonica* (alignment length > 1000, alignment identity > 90), then RepeatModeler (http://www.repeatmasker.org/) was used to create repeat sequences database based on above matching sequences (RMBlast), and RepeatMasker was used for identify TEs in subgenomes. Subsequently, the substitutions of these identified TEs were calculated with calcDivergenceFromAlign.pl (a script in RepeatMasker). Divergence was obtained by comparing TEs between *S. babylonica* and the repeat sequences database obtained above. Finally, the ggplot2 (88) R package was used to print divergence values using GAM (Generalized Additive Model) to fit the curves. Two time-points of divergence and merger between two subgenomes corresponded to TE divergence rates of 18% and 2.5% in Aa vs. Ba, as well as 23% and 1.5% in Aa vs. Bb (Fig S6). The divergence time of *Salix* and *Vetrix* was 42.13 Mya according to the result of phylogenetic analysis above. Therefore, the allotetraploidization time was estimated: 42.13 Ma×2.5/18 ≈ 5.85 Mya and 42.13 Ma×1.5/23 ≈ 2.75 Mya.

### Identification of the sex-linked regions of *Salix dunnii* and weeping willow

Sequence reads of weeping willow and *S. dunnii* were filtered and trimmed by fastp (89) with parameters “--cut_by_quality3 --cut_by_quality5 --n_base_limit 0 --length_required 60 --correction”. The KMC 3.1.1 (90) was used to count the *31*-mer of each clean dataset of weeping willow. The *k*-mers with less than 30% missing and average count > 1 in each sex group were obtained. The *k*-mers in all 40 individuals with minor allele (*k*-mer absent or present) frequency <0.1 were discarded. The filtered *k*-mers were mapped to the genome (76 chromosomes) of weeping willow using Bowtie (91). *K-*mers with coverage of 0 in all of 20 female individuals were defined as male-specific *k-*mers, while *k-*mers with coverage of 0 in 20 male individuals were defined as female-specific *k-*mers. We expect significantly male (male heterogamety) or female (female heterogamety) specific *k*-mers on sex-liked region of sex chromosome(s).

We employed the CQ method (37) to detect sex chromosomes of the two willows. We made combined female and male clean read data sets of the two willows, respectively. The cq-calculate.pl software (37) was used to calculate the CQ for each 50-kb nonoverlapping window of the genomes. For male heterogamety, the CQ is the normalized ratio of female to male alignments to a given reference sequence, and CQ value close to 2 (for XX/XY, 1.33 for XXXX/XXXY) in windows in the X-linked region and to zero in windows in the Y-linked region. For female heterogamety, the CQ is the normalized ratio of male to female alignments, and CQ value close to 2 (for ZW/ZZ, 1.33 for ZZZW/ZZZZ) in windows in Z-linked region and to zero in windows in the W-linked region.

Clean reads of *S. dunnii* were aligned to its genome (two haplotypes) using the BWA-MEM algorithm from bwa 0.7.12 (92, 93) with default parameters. Samtools 0.1.19 (94) was used to extract primary alignments, sort, and merge the mapped data. PCR replicates were filtered using sambamba 0.7.1 (95). The variants were called and filtered using Genome Analysis Toolkit v. 4.1.8.1 and VCFtools 0.1.16 (96). Hard filtering of the SNP calls was carried out with “QD< 2.0, FS > 60.0, MQ< 40.0, MQRankSum<−12.5, ReadPosRankSum <−8.0, SOR > 3.0”. Only biallelic sites were kept for subsequent filtering. The sites with coverage greater than twice the mean depth at all variant sites across all samples were discarded. Genotypes with depth < four were treated as missing, sites with >10% missing data were filtered out, and sites with minor allele frequency < 0.05 were discarded. We used VCFtools to calculate weighted *F*_ST_ values between the sexes using the (97) estimator with 100-kb windows and 10-kb steps. The Changepoint package (98) was used to detect the boundaries of the sex-linked regions based on *F*_ST_ and CQ values between the sexes of *S. dunnii* and CQ values and sex specific *k*-mer counts of weeping willow, respectively. Furthermore, we used the Python version of MCScan (99) to analyze chromosome collinearity between the protein coding sequences detected in 7 and 15 chromosomes of *S. dunnii* and weeping willow and their homologous autosomes, to detect possible inversions in sex chromosomes. The “--cscore=.99” was used to obtain reciprocal best hit (RBH) orthologues for the collinearity analysis. We then combined all the previously analyses to detect the sex-linked regions.

### *ARR17* identification and phylogeny

In order to obtain *ARR17*-like gene duplicates in target species, we used BLASTN to blast *ARR17*-like gene (Potri.019G133600 (32)) against the *S. arbutifolia* (34), *S. dunnii*, and *S. babylonica* genomes with parameters “-evalue 1e-5 -word_size 8”. *ARR17*-like gene includes five CDSs, so only the genes including all the CDSs were classified as intact *ARR17*-like gene duplicates, while the sequences with fewer than five CDSs were regarded as partial *ARR17*-like gene duplicates (33). We combined different CDSs of the same *ARR17*-like gene to construct the phylogeny. These CDSs were aligned with MAFFT (81), then we constructed a phylogenetic tree with IQ-TREE.

### Ancient sex-linked genes identification

We used the OrthoFinder (80) to identify single copy genes in sex-linked regions of 15W and 15Z of *S. babylonica* and *S. purpurea*, 15X and 15Y of *S. arbutifolia*, and chromosome 15a of *S. dunnii*. We then extracted homologous genes, that diverged between ancestral X and Y in ancestors of *Vetrix* clade and close to partial *ARR17*-like gene duplicates, proposed by (34). We obtained one ancient single-copy homologous genes, then used MAFFT and IQ-TREE to align and reconstruct the phylogenetic tree using *S. dunnii* as outgroup.

### Sex-linked region features and gene loss

We calculated content of gene and repeat sequences (total repeat, TE, Gypsy, Copia) in PAR and sex-linked region among *S. babylonica* (15W, 15Z) and *S. dunnii* (7X, 7Y) (Table S12). And we also used 100kb window to show changes between sex-linked regions and the PAR (Fig. S17).

We used 15a in *S. dunnii* as reference to identified the protein-coding gene loss in 15W, 15Z, 15X and 15Y, and used 7Ba in *S. babylonica* as reference to identified the gene loss in 7X and 7Y. Firstly, we identified shared genes in W-Z and X-Y, and specific genes in W, Z, X, and Y. For specific genes in X-SLR/W-SLR, there is no homologous gene in Y-SLR/Z-SLR. Similarly, there is no homologous gene in X-SLR/W-SLR for specific genes in Y-SLR/Z-SLR. Then we used these specific genes to determine gene loss situation among them. The degradation rate of W-SLR = W-SLR loss / (W-SLR loss + Z-SLR loss + their shared genes), and the degradation rated of Z-SLR = Z-SLR loss / (W-SLR loss + Z-SLR loss + their shared genes). Similarly, we obtained the degradation rate of X-SLR and Y-SLR. We also obtained the relevant data of PARs.

### Gene expression analyses

We calculated the gene expression among female and male catkins of *S. babylonica*, *S. arbutifolia*, and *S. dunnii*, respectively. The RNA datasets of *S. arbutifolia* are from (34). Each sex and individual contained three independent biological replicates (Table S16, (34)). After filtering, clean transcript reads from each sample were mapped to the their own genome with HISAT2 (100). We used a haplotype genome assembly and sex chromosomes (15X and 15Y or 7X and 7Y) as reference genome in *S. arbutifolia* and *S. dunnii*, and Aa and Ba haplotypes genome assembly and sex chromosomes (15W and 15Z) as reference genome in *S. babylonica*. The number of reads mapping to each gene was calculated using featureCounts (101). Then we converted these read counts to TPM (transcripts per million reads). After filtering out unexpressed genes (counts=0 in all samples, excluding non-mRNA), then we used the expression levels of the sex-linked regions in male, female and their autosomes to identify the dosage compensation pattern (102). For complete compensation patterns, the expression levels of male, female and their autosomes are similar.

## Acknowledgements

This study was financially supported by the National Natural Science Foundation of China (grant no. 32171813) and Special Fund for Scientific Research of Shanghai Landscaping & City Appearance Administrative Bureau (grant nos. G232403 & G242417). We are grateful to Si-Wen Zeng for sampling. We are indebted to James H. Leebens-Mack, Zhenyang Liao, Zhiqing Xue, Guangnan Gong for their kind help during preparation of our paper.

## Data Availability Statement

The genome assembly sequences have been deposited in the Genome Warehouse (GWH), which is available at https://bigd.big.ac.cn/gwh, at the National Genomics Data Center (NGDC) with the BioProject number: PRJCA016000. The sequencing datasets have been deposited in NCBI under the BioProject accession numbers PRJNA882493, PRJNA1020583, & PRJNA1020619.

## References

1. B. J. Evans, R. Alexander Pyron, J. J. Wiens, “Polyploidization and sex chromosome evolution in amphibians” in Polyploidy and Genome Evolution, (Springer, 2012), pp. 385–410.

2. K. Van de Peer, Yves and Mizrachi, Eshchar and Marchal, The evolutionary significance of polyploidy. Nat. Rev. Genet. 18, 411–424 (2017).

3. H. J. Muller, Why polyploidy is rarer in animals than in plants. Am. Nat. 59, 346–353 (1925).

4. L. He, E. Hörandl, Does polyploidy inhibit sex chromosome evolution in angiosperms? Front. Plant Sci. 13, 976765 (2022).

5. B. K. Mable, M. A. Alexandrou, M. I. Taylor, Genome duplication in amphibians and fish: an extended synthesis. J. Zool. 284, 151–182 (2011).

6. J. Ramsey, D. W. Schemske, Pathways, mechanisms, and rates of polyploid formation in flowering plants. Annu. Rev. Ecol. Syst. 29, 467–501 (1998).

7. L. He, et al., Evolutionary origin and establishment of a dioecious diploid-tetraploid complex. Mol. Ecol. 32, 2732–2749 (2023).

8. M. Stöck, et al., Sex chromosomes in meiotic, hemiclonal, clonal and polyploid hybrid vertebrates: along the ‘extended speciation continuum’’.’ Philos. Trans. R. Soc. B 376, 20200103 (2021).

9. S. Gulyaev, et al., The phylogeny of Salix revealed by whole genome re-sequencing suggests different sex-determination systems in major groups of the genus. Ann. Bot. 129, 485–498 (2022).

10. S. P. Otto, J. Whitton, Polyploid incidence and evolution. Annu. Rev. Genet. 34, 401–437 (2000).

11. M. F. Scott, M. M. Osmond, S. P. Otto, Haploid selection, sex ratio bias, and transitions between sex-determining systems. PLoS Biol. 16, e2005609 (2018).

12. S. Mawaribuchi, et al., Sex chromosome differentiation and the W-and Z-specific loci in *Xenopus laevis*. Dev. Biol. 426, 393–400 (2017).

13. F. M. Robertson, et al., Lineage-specific rediploidization is a mechanism to explain time-lags between genome duplication and evolutionary diversification. Genome Biol. 18, 1–14 (2017).

14. R. R. Gates, Polyploidy and sex chromosomes. Nature 117, 234 (1926).

15. N. Cunado, et al., The evolution of sex chromosomes in the genus *Rumex* (Polygonaceae): identification of a new species with heteromorphic sex chromosomes. Chromosom. Res. 15, 825–833 (2007).

16. J. K. Abbott, A. K. Nordén, B. Hansson, Sex chromosome evolution: Historical insights and future perspectives. Proc. R. Soc. B Biol. Sci. 284 (2017).

17. D. Charlesworth, B. Charlesworth, G. Marais, Steps in the evolution of heteromorphic sex chromosomes. Heredity (Edinb). 95, 118–128 (2005).

18. T. Lenormand, D. Roze, Y recombination arrest and degeneration in the absence of sexual dimorphism. Science (80-.). 375, 663–666 (2022).

19. J. Yue, et al., The origin and evolution of sex chromosomes, revealed by sequencing of the *Silene latifolia* female genome. Curr. Biol. (2023).

20. Q. Zhou, et al., Complex evolutionary trajectories of sex chromosomes across bird taxa. Science (80-.). 346 (2014).

21. B. Charlesworth, P. Sniegowski, W. Stephan, The evolutionary dynamics of repetitive DNA in eukaryotes. Nature 371, 215–220 (1994).

22. H. Ellegren, Sex-chromosome evolution: recent progress and the influence of male and female heterogamety. Nat. Rev. Genet. 12, 157–166 (2011).

23. A. Muyle, et al., Rapid de novo evolution of X chromosome dosage compensation in *Silene latifolia*, a plant with young sex chromosomes. PLoS Biol. 10, e1001308 (2012).

24. B. Charlesworth, Model for evolution of Y chromosomes and dosage compensation. Proc. Natl. Acad. Sci. 75, 5618–5622 (1978).

25. J. E. Mank, Sex chromosome dosage compensation: definitely not for everyone. Trends Genet. 29, 677–683 (2013).

26. A. Muyle, R. Shearn, G. A. B. Marais, The evolution of sex chromosomes and dosage compensation in plants. Genome Biol. Evol. 9, 627–645 (2017).

27. S. Ohno, Sex chromosomes and sex linked genes, S. L. Labhart A, Mann T, Ed. (Springer, Heidelberg, 1967) 10.1111/j.1439-0272.1973.tb00879.x.

28. Z. Fang, S. D. Zhao, A. K. Skvortsov, Salicaceae. Flora of China 4, 139–274 (1999).

29. J. G. Isebrands, J. Richardson, Poplars and willows: trees for society and the environment (CABI, 2014).

30. Y. Wang, et al., Correction: The male-heterogametic sex determination system on chromosome 15 of *Salix triandra* and *Salix arbutifolia* reveals ancestral male heterogamety and subsequent turnover events in the genus *Salix*. Heredity (Edinb). 130, 177 (2023).

31. S. C. Teng, H. Yu, Propagation of weeping willow from seed. Bot. Bull. Acad. Sin. 2, 131–132 (1948).

32. N. A. Müller, et al., A single gene underlies the dynamic evolution of poplar sex determination. Nat. plants 6, 630–637 (2020).

33. D. Wang, et al., Repeated turnovers keep sex chromosomes young in willows. Genome Biol. 23, 1–23 (2022).

34. Y. Wang, et al., Gap-free X and Y chromosomes of *Salix arbutifolia* reveal an evolutionary change from male to female heterogamety in willows, without a change in the sex-determining region. bioRxiv (2023) 10.1101/2023.10.11.561967.

35. L. He, et al., Chromosome-scale assembly of the genome of *Salix dunnii* reveals a male-heterogametic sex determination system on chromosome 7. Mol. Ecol. Resour. 21, 1966–1982 (2021).

36. J. Wu, et al., Phylogeny of *Salix* subgenus *Salix* sl (Salicaceae): delimitation, biogeography, and reticulate evolution. BMC Evol. Biol. 15, 1–13 (2015).

37. A. B. Hall, et al., Six novel Y chromosome genes in *Anophelesmosquitoes* discovered by independently sequencing males and females. BMC Genomics 14, 1–13 (2013).

38. J. R. Harlan, J. M. J. deWet, On Ö. Winge and a prayer: the origins of polyploidy. Bot. Rev. 41, 361–390 (1975).

39. A. E. Wright, R. Dean, F. Zimmer, J. E. Mank, How to make a sex chromosome. Nat. Commun. 7, 12087 (2016).

40. A. S. T. Papadopulos, M. Chester, K. Ridout, D. A. Filatov, Rapid Y degeneration and dosage compensation in plant sex chromosomes. Proc. Natl. Acad. Sci. 112, 13021–13026 (2015).

41. R. Ming, A. Bendahmane, S. S. Renner, Sex chromosomes in land plants. Annu. Rev. Plant Biol. 62, 485–514 (2011).

42. D. Prentout, et al., An efficient RNA-seq-based segregation analysis identifies the sex chromosomes of *Cannabis sativa*. Genome Res. 30, 164–172 (2020).

43. D. W. Bellott, et al., Convergent evolution of chicken Z and human X chromosomes by expansion and gene acquisition. Nature 466, 612–616 (2010).

44. T. Akagi, I. M. Henry, T. Kawai, L. Comai, R. Tao, Epigenetic regulation of the sex determination gene *MeGI* in polyploid persimmon. Plant Cell 28, 2905–2915 (2016).

45. J. F. Gerchen, P. Veltsos, J. R. Pannell, Recurrent allopolyploidization, Y-chromosome introgression and the evolution of sexual systems in the plant genus *Mercurialis*. Philos. Trans. R. Soc. B 377, 20210224 (2022).

46. C. M. S. Cauret, S. M. E. Mortimer, M. C. Roberti, T.-L. Ashman, A. Liston, Chromosome-scale assembly with a phased sex-determining region resolves features of early Z and W chromosome differentiation in a wild octoploid strawberry. G3 12, jkac139 (2022).

47. G. Hillman, R. Hedges, A. Moore, S. Colledge, P. Pettitt, New evidence of Lateglacial cereal cultivation at Abu Hureyra on the Euphrates. The Holocene 11, 383–393 (2001).

48. P. Almeida, et al., Genome assembly of the basket willow, *Salix viminalis*, reveals earliest stages of sex chromosome expansion. BMC Biol. 18, 1–18 (2020).

49. R. Zhou, et al., A willow sex chromosome reveals convergent evolution of complex palindromic repeats. Genome Biol. 21, 1–19 (2020).

50. J. Wang, et al., Sequencing papaya X and Yh chromosomes reveals molecular basis of incipient sex chromosome evolution. Proc. Natl. Acad. Sci. 109, 13710–13715 (2012).

51. J. Hough, J. D. Hollister, W. Wang, S. C. H. Barrett, S. I. Wright, Genetic degeneration of old and young Y chromosomes in the flowering plant *Rumex hastatulus*. Proc. Natl. Acad. Sci. 111, 7713–7718 (2014).

52. D. Bachtrog, Y-chromosome evolution: emerging insights into processes of Y-chromosome degeneration. Nat. Rev. Genet. 14, 113–124 (2013).

53. J. Doležel, J. Greilhuber, J. Suda, Estimation of nuclear DNA content in plants using flow cytometry. Nat. Protoc. 2, 2233–2244 (2007).

54. G. Marçais, C. Kingsford, A fast, lock-free approach for efficient parallel counting of occurrences of k-mers. Bioinformatics 27, 764–770 (2011).

55. T. R. Ranallo-Benavidez, K. S. Jaron, M. C. Schatz, GenomeScope 2.0 and Smudgeplot for reference-free profiling of polyploid genomes. Nat. Commun. 11, 1–10 (2020).

56. J.-M. Belton, et al., Hi--C: a comprehensive technique to capture the conformation of genomes. Methods 58, 268–276 (2012).

57. H. Cheng, G. T. Concepcion, X. Feng, H. Zhang, H. Li, Haplotype-resolved de novo assembly using phased assembly graphs with hifiasm. Nat. Methods 18, 170–175 (2021).

58. N. C. Durand, et al., Juicer provides a one-click system for analyzing loop-resolution Hi-C experiments. Cell Syst. 3, 95–98 (2016).

59. O. Dudchenko, et al., De novo assembly of the Aedes aegypti genome using Hi-C yields chromosome-length scaffolds. Science (80-.). 356, 92–95 (2017).

60. N. C. Durand, et al., Juicebox provides a visualization system for Hi-C contact maps with unlimited zoom. Cell Syst. 3, 99–101 (2016).

61. G.-C. Xu, et al., LR_Gapcloser: a tiling path-based gap closer that uses long reads to complete genome assembly. Gigascience 8, giy157 (2019).

62. J. Hu, J. Fan, Z. Sun, S. Liu, NextPolish: a fast and efficient genome polishing tool for long-read assembly. Bioinformatics 36, 2253–2255 (2020).

63. J.-J. Jin, et al., GetOrganelle: a fast and versatile toolkit for accurate de novo assembly of organelle genomes. Genome Biol. 21, 1–31 (2020).

64. L. P. Pryszcz, T. Gabaldón, Redundans: an assembly pipeline for highly heterozygous genomes. Nucleic Acids Res. 44, e113--e113 (2016).

65. M. G. Grabherr, et al., Full-length transcriptome assembly from RNA-Seq data without a reference genome. Nat. Biotechnol. 29, 644–652 (2011).

66. D. Kim, B. Langmead, S. L. Salzberg, HISAT: a fast spliced aligner with low memory requirements. Nat. Methods 12, 357–360 (2015).

67. M. Pertea, et al., StringTie enables improved reconstruction of a transcriptome from RNA-seq reads. Nat. Biotechnol. 33, 290–295 (2015).

68. L. Fu, B. Niu, Z. Zhu, S. Wu, W. Li, CD-HIT: accelerated for clustering the next-generation sequencing data. Bioinformatics 28, 3150–3152 (2012).

69. B. J. Haas, et al., Improving the *Arabidopsis* genome annotation using maximal transcript alignment assemblies. Nucleic Acids Res. 31, 5654–5666 (2003).

70. M. Stanke, M. Diekhans, R. Baertsch, D. Haussler, Using native and syntenically mapped cDNA alignments to improve de novo gene finding. Bioinformatics 24, 637–644 (2008).

71. B. L. Cantarel, et al., MAKER: an easy-to-use annotation pipeline designed for emerging model organism genomes. Genome Res. 18, 188–196 (2008).

72. M. Stanke, et al., AUGUSTUS: ab initio prediction of alternative transcripts. Nucleic Acids Res. 34, W435--W439 (2006).

73. G. S. C. Slater, E. Birney, Automated generation of heuristics for biological sequence comparison. BMC Bioinformatics 6, 1–11 (2005).

74. B. J. Haas, et al., Automated eukaryotic gene structure annotation using EVidenceModeler and the Program to Assemble Spliced Alignments. Genome Biol. 9, 1–22 (2008).

75. R.-G. Zhang, et al., TEsorter: an accurate and fast method to classify LTR-retrotransposons in plant genomes. Hortic. Res. 9, uhac017 (2022).

76. T. M. Lowe, S. R. Eddy, tRNAscan-SE: a program for improved detection of transfer RNA genes in genomic sequence. Nucleic Acids Res. 25, 955–964 (1997).

77. J. Huerta-Cepas, et al., Fast genome-wide functional annotation through orthology assignment by eggNOG-mapper. Mol. Biol. Evol. 34, 2115–2122 (2017).

78. P. Jones, et al., InterProScan 5: genome-scale protein function classification. Bioinformatics 30, 1236–1240 (2014).

79. S. Ou, et al., Benchmarking transposable element annotation methods for creation of a streamlined, comprehensive pipeline. Genome Biol. 20, 1–18 (2019).

80. D. M. Emms, S. Kelly, OrthoFinder: phylogenetic orthology inference for comparative genomics. Genome Biol. 20, 1–14 (2019).

81. K. Katoh, D. M. Standley, MAFFT multiple sequence alignment software version 7: improvements in performance and usability. Mol. Biol. Evol. 30, 772–780 (2013).

82. B. Q. Minh, et al., IQ-TREE 2: New Models and Efficient Methods for Phylogenetic Inference in the Genomic Era. Mol. Biol. Evol. 37, 1530–1534 (2020).

83. C. Zhang, M. Rabiee, E. Sayyari, S. Mirarab, ASTRAL-III: polynomial time species tree reconstruction from partially resolved gene trees. BMC Bioinformatics 19, 15–30 (2018).

84. Z. Yang, PAML 4: phylogenetic analysis by maximum likelihood. Mol. Biol. Evol. 24, 1586–1591 (2007).

85. G. Bi, Y. Mao, Q. Xing, M. Cao, HomBlocks: a multiple-alignment construction pipeline for organelle phylogenomics based on locally collinear block searching. Genomics 110, 18–22 (2018).

86. P. Xu, et al., The allotetraploid origin and asymmetrical genome evolution of the common carp Cyprinus carpio. Nat. Commun. 10, 1–11 (2019).

87. A. L. Delcher, S. L. Salzberg, A. M. Phillippy, Using MUMmer to identify similar regions in large sequence sets. Curr. Protoc. Bioinforma., 10–13 (2003).

88. H. Wickham, ggplot2: Elegant Graphics for Data Analysis (Springer-Verlag New York, 2016).

89. S. Chen, Y. Zhou, Y. Chen, J. Gu, fastp: an ultra-fast all-in-one FASTQ preprocessor. Bioinformatics 34, i884--i890 (2018).

90. M. Kokot, M. Długosz, S. Deorowicz, KMC 3: counting and manipulating k-mer statistics. Bioinformatics 33, 2759–2761 (2017).

91. B. Langmead, C. Trapnell, M. Pop, S. L. Salzberg, Ultrafast and memory-efficient alignment of short DNA sequences to the human genome. Genome Biol. 10, 1–10 (2009).

92. H. Li, Aligning sequence reads, clone sequences and assembly contigs with BWA-MEM. arXiv Prepr. arXiv1303.3997 (2013).

93. H. Li, R. Durbin, Fast and accurate short read alignment with Burrows--Wheeler transform. bioinformatics 25, 1754–1760 (2009).

94. H. Li, et al., The sequence alignment/map format and SAMtools. bioinformatics 25, 2078–2079 (2009).

95. A. Tarasov, A. J. Vilella, E. Cuppen, I. J. Nijman, P. Prins, Sambamba: fast processing of NGS alignment formats. Bioinformatics 31, 2032–2034 (2015).

96. P. Danecek, et al., The variant call format and VCFtools. Bioinformatics 27, 2156–2158 (2011).

97. B. S. Weir, C. C. Cockerham, Estimating F-statistics for the analysis of population structure. Evolution (N. Y)., 1358–1370 (1984).

98. R. Killick, I. Eckley, changepoint: An R package for changepoint analysis. J. Stat. Softw. 58, 1–19 (2014).

99. H. Tang, et al., Unraveling ancient hexaploidy through multiply-aligned angiosperm gene maps. Genome Res. 18, 1944–1954 (2008).

100. D. Kim, J. M. Paggi, C. Park, C. Bennett, S. L. Salzberg, Graph-based genome alignment and genotyping with HISAT2 and HISAT-genotype. Nat. Biotechnol. 37, 907–915 (2019).

101. Y. Liao, G. K. Smyth, W. Shi, featureCounts: an efficient general purpose program for assigning sequence reads to genomic features. Bioinformatics 30, 923–930 (2014).

102. I. Darolti, et al., Extreme heterogeneity in sex chromosome differentiation and dosage compensation in livebearers. Proc. Natl. Acad. Sci. 116, 19031–19036 (2019).

